# Global determinants of navigation ability

**DOI:** 10.1101/188870

**Authors:** A. Coutrot, R. Silva, E. Manley, W. de Cothi, S. Sami, V. D. Bohbot, J. M. Wiener, C. Hölscher, R.C. Dalton, M. Hornberger, H. J. Spiers

**Author notes:** and. These authors contributed equally to this work.

## Abstract

Countries vary in their geographical and cultural properties. Only a few studies have explored how such variations influence how humans navigate or reason about space [1–7]. We predicted that these variations impact human cognition, resulting in an organized spatial distribution of cognition at a planetary-wide scale. To test this hypothesis we developed a mobile-app-based cognitive task, measuring non-verbal spatial navigation ability in more than 2.5 million people, sampling populations in every nation state. We focused on spatial navigation due to its universal requirement across cultures. Using a clustering approach, we find that navigation ability is clustered into five distinct, yet geographically related, groups of countries. Specifically, the economic wealth of a nation was predictive of the average navigation ability of its inhabitants, and gender inequality was predictive of the size of performance difference between males and females. Thus, cognitive abilities, at least for spatial navigation, are clustered according to economic wealth and gender inequalities globally, which has significant implications for cross-cultural studies and multi-centre clinical trials using cognitive testing.

---

In the *Origin of Certain Instincts*, Charles Darwin alludes to the differences in navigation abilities between cultures, comparing the natives of Siberia to his European companion: “[Von Wrangell], an experienced surveyor, and using a compass, failed to do that which these savages easily effected” [8]. Since then only a few studies have explored how people from different cultures differ in how they navigate or reason about space [1–7]. Here, we explored how spatial navigation abilities vary on a global scale. To achieve this we devised a mobile video game designed to measure human spatial navigation ability through gameplay - Sea Hero Quest (SHQ). The game involves navigating a boat in search of sea creatures in order to photograph them (Figure 1 and Video S1). It features two main tasks: wayfinding and path integration. In wayfinding levels, players are initially presented with a map indicating start location and the location of several checkpoints to find in a set order (Figure 1A-C and Figure S2). Wayfinding task requires quite elaborate processing including interpretation of a map, planning a multi-stop route, memory of the route, monitoring progress along the route and updating of route plan, transformation of birds-eye perspective to an egocentric perspective needed for navigation [9]. In path integration levels, participants navigate along a river with bends to find a flare gun and then choose which three directions is the correct direction back to the starting point along the Euclidean (Figure 1D and Figure S3). During path integration, one integrates perceived ego motion during travel to update one’s position and orientation. It is a more basic (and evolutionary highly conserved) navigation mechanism, which typically only requires working memory processes [10, 11]. Together, wayfinding and path integration capture a wide range of the abilities and processes that are required for everyday successful navigation. 2,512,123 people between 18 and 99 years old from all 195 countries in the world downloaded and played the game (details in Table S1). 57.6% of the participants provided demographics of their age, gender and nationality (Figure S4). To provide a reliable estimate of spatial navigation ability we examined the data only from those subjects who had completed a minimum of 9 levels of the game (see Methods). This resulted in 558,143 participants from 57 countries that were included in our analysis (Table S1).

**Figure 1.**
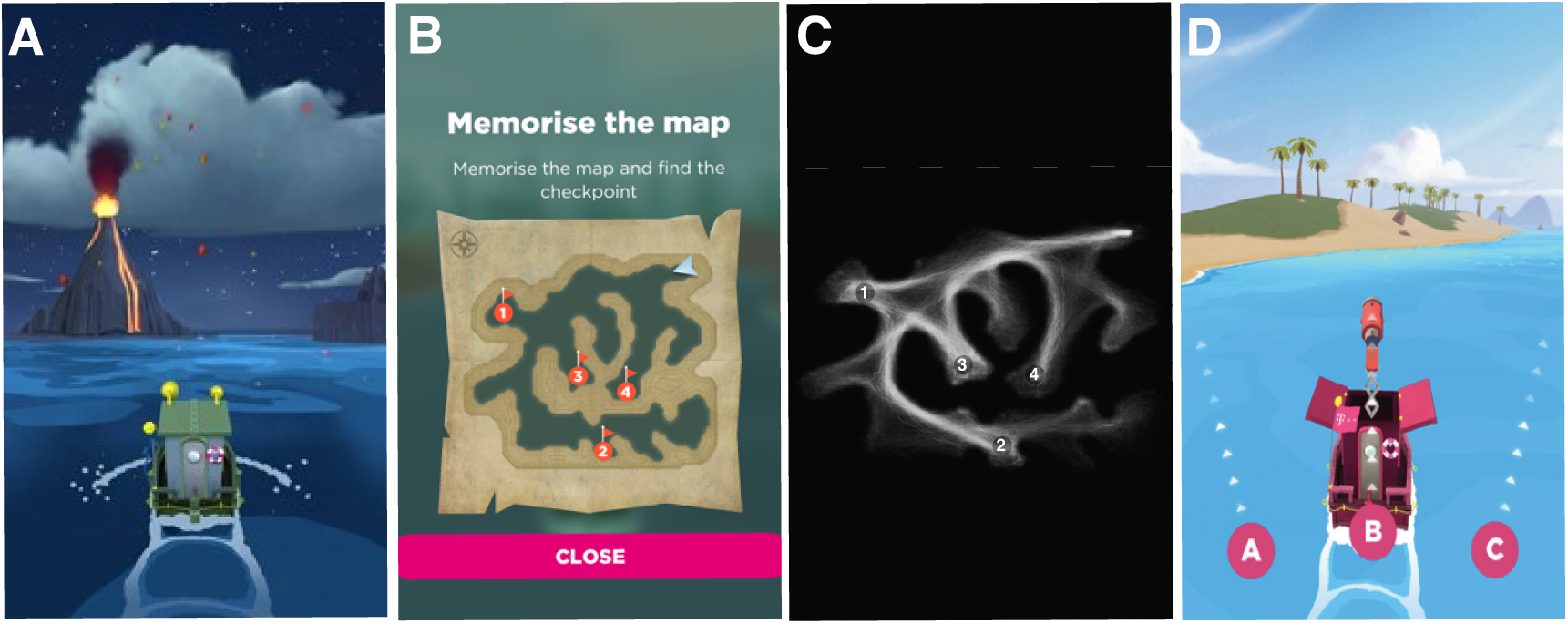
Tasks design. (A-B) Wayfinding task: a map of the level featuring the ordered set of checkpoints to reach is presented and disappears when the game starts. (C) Superposition of 1000 individual trajectories randomly sampled from level 32. (D) Path integration task: after navigating the level, participants must shoot a flare back to the starting point.

To quantify spatial abilities, three measures were computed: trajectory length, duration of navigation epoch for wayfinding; overall accuracy for path integration. We defined an overall performance (OP) metric as the first component of a Principal Component Analysis across these three measures. OP captures different aspects of navigation abilities, providing a general performance measure, which reflects overall navigation abilities covering a range of tasks. We considered that video games experience might bias performance, with players familiar with similar games having an advantage. Therefore we normalized OP with performance on the first two tutorial wayfinding levels, where no navigation skill was required (Figure S2d,e: goals are visible from the starting point.). We discuss and control for other potential biases such as the unavailability of SHQ in some languages, ‘fake’ demographics, and the virtual nature of the task in Supplementary Discussion and Figure S8 to S12.

## Results and Discussion

Across all countries we observed a similar pattern of decline in ability with age and a male advantage between 19 and 60 years old (Figure 2E, see Figure S13 for plots from example nations). This result held true in all tested countries after correcting for differences in age and gender distributions. We computed a multiple linear regression to predict OP based on gender and age. Gender estimate had the same sign in every country, ranging from 0.30 to 1.59, *M* = 0.96, 95%*IC* = [0.89,1.04], Cohen’s *d* ranging from 0.09 to 0.48, *M* = 0.29,95%*IC* = [0.27,0.31], Extended Figure S14. While a number of previous studies have examined the size of gender differences in cognitive abilities across countries, the underlying causes of such variation are still debated. Advocates of the *gender stratification hypothesis* argue that gender differences are more pronounced in countries with less equity [12, 13]. Data from Programme for International Student Assessment (PISA) that reports on more than 250,000 15-year-old students from 40 countries show that the gender gap in math scores disappears in countries with a more gender-equal culture [14, 15]. By contrast, other studies link gender differences more to evolved sex-linked dispositions and environmental affordances [16]. For example, difference in mental rotation and line angle judgment performance in more than 200,000 men and women from 53 nations remained even when controlling for gender equality [17]. Here, we report a positive correlation between the magnitude of gender differences and gender inequalities assessed by the World Economic Forum’s Gender Gap Index (GGI), which reflects economic and political opportunities, education, and well-being for women (Figure 2D, *ρ* = 0.52, *p* < .001). We computed a multiple linear regression to predict gender estimates based on GDP and GGI. Both GGI (*t*(52) = −2.45, *p* = 0.01) and GDP per capita (*t*(52) = −2.87, *p* = 0.006) significantly predicted Countries’ gender estimates. This suggests that the gender effect is not just related to countries’ wealth, but also to the improvement of the role of women in society.

**Figure 2.**
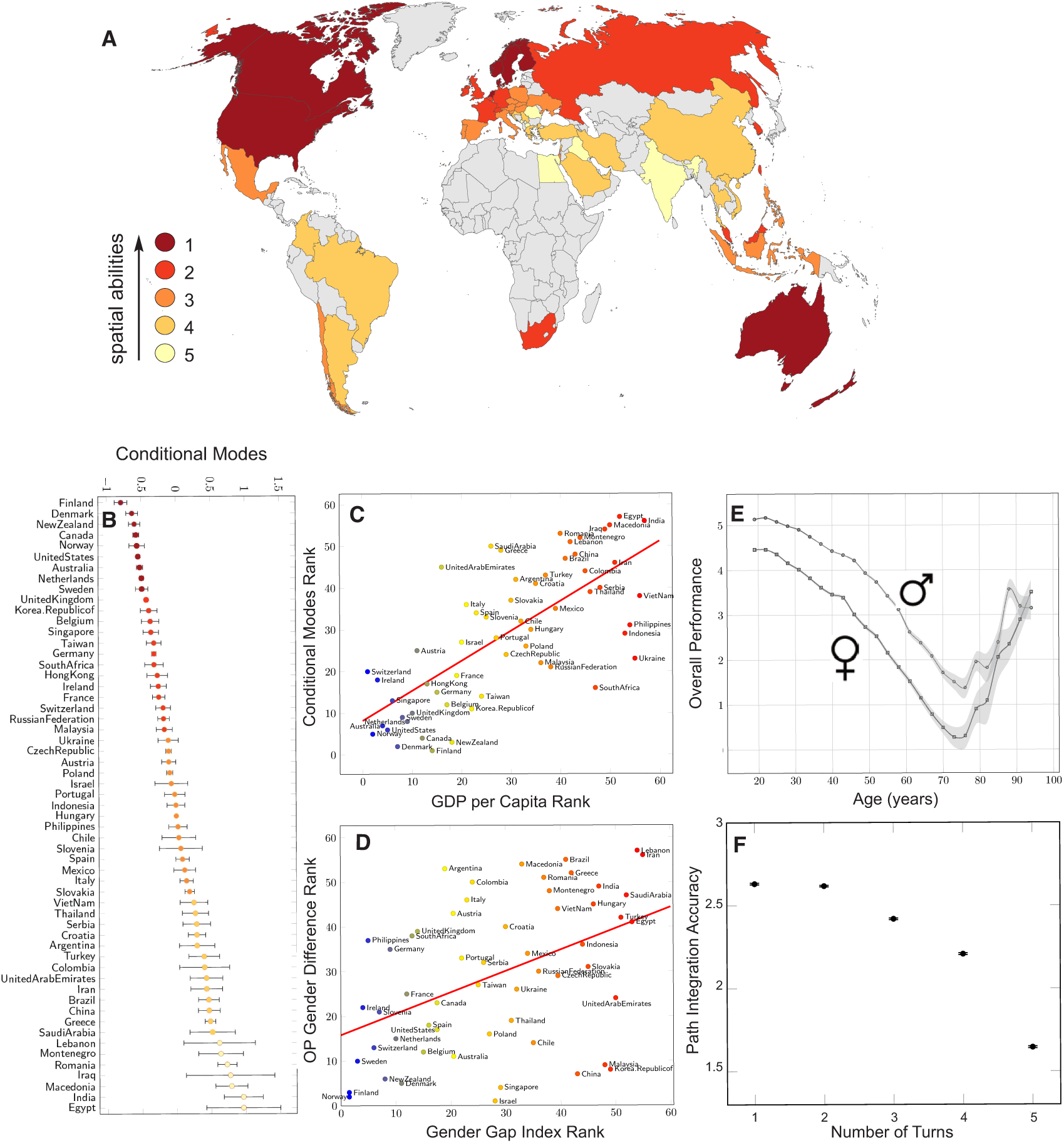
Spatial ability distribution across age, gender, and nations. (A-B) Five world clusters of people with similar overall performance (OP). OP is the first component of a Principal Component Analysis (PCA) across duration, trajectory length and flare accuracy corrected for video gaming skill. Conditional Modes (CM) are the difference between the global average predicted response and the response predicted for a particular country (the lower the better). (C) Correlation between country performance (CM) and GDP per capita (r=0.69, p¡0.001). (D) Correlation between gender estimates from a multiple linear regression (OP age + gender) and Gender Gap Index (r=0.52, p¡0.001). (E) Evolution of OP across age and gender. Data points correspond to the average OP within 3-year windows. Error bars correspond to standard errors. (F) Path integration accuracy vs. path complexity. Path integration accuracy is defined as the average number of stars obtained by participants (1, 2 or 3 stars). With increasing complexity: level 14 (1 turn), level 34 (2 turns), level 54 (3 turns), level 44 (4 turns) and level 74 (five turns). This plot includes participants that completed all five levels (N = 19,038). For more details see Figure S10. Error bars correspond to standard errors.

The age related decline in navigation abilities - OP decreases in a linear fashion between 19 and 60 years-old (Figure 2E) - held true in all tested countries, age estimates ranging from −0.092 to −0.022 per year, *M* = −0.059, 95%*IC* = [−0.063,−0.055], Figure S13. Our observed early decline in performance mirrors the decline in ‘fluid intelligence’ components such as reasoning and working memory, which generally occurs in healthy adults [18–20]. Our observed performance increment after 70 years of age was not predicted from the past literature and is consistent with a selection bias that those older participants willing to participate in online testing are likely to have greater cognitive skills. At the individual level performance should continue to decline, as demonstrated in prior studies of navigation in elderly humans [21–24].

To our knowledge no prior large-scale studies have quantified the impact of nationality on a cognitive task. To assess the impact of nationality on spatial navigation we fit a multilevel model for OP, with fixed effects for age and gender and random effect for nationality: *OP* ~ *age* + *gender* + (1|*nationality*). Multilevel - or mixed-effects - models take into account the shared variance between neighboring countries by modeling the covariance between them. When we compare our multi-level model with a single-level model including only age and gender, nationality has a significant impact on OP (χ_2_(1) = 6413.8, *p* < 0.001). The variance partition coefficient (VPC) indicates that 1.7% of the variance in performance can be attributed to differences between nationalities. Figure 2B represents countries ranked according to their conditional modes (CM), that is the difference between the global average predicted response in performance and the response predicted for a particular country. A reasonable assumption is that while OP differs around the world, it follows a relatively smooth uniform distribution with some countries populations performing well and other performing less well on average. An alternative possibility is that countries are grouped by similar cognitive strategies, and that some countries will tend to behave more similarly. Indeed, country-level might not be the optimal scale to work at, since many social and geographical traits know no borders. To the best of our knowledge this hypothesis has never been tested with data from cognitive tests. To address this we pooled countries with similar CM into *k* clusters via the optimal 1D *k*-means algorithm [25]. We defined the optimal *k* as the one maximizing VPC. This was achieved for *k* = 5, VPC = 2.6%, Figure 2A and Figure S8. Thus, spatial navigation ability appears to be clustered. Importantly, this clustering is distinct from GDP per Capita and video gaming skill distributions across countries (Figure S8). We downsampled the data to equate video gaming skill in our population and found a ranking and clustering nearly identical to the one with the full dataset (Pearson’s correlation *ρ* = 0.99, *p* < 0.001, Figure S7 and 9).

The clustering of navigation abilities is not geographically random. Indeed, countries’ CM were correlated with Gross Domestic Product (GDP) per capita (Figure 2C, Pearson’s correlation *ρ* = 0.69, *p* < 0.001). This can be explained by different variables highly correlated with GDP associated with better spatial abilities, such as level of education [26] - particularly in science [27,28] - or ability to travel [29]. Figure S16 shows a positive correlation between countries’ CM and average scores at PISA 2015 (Pearson’s correlation *ρ* = 0.73, *p* < 0.001).

While GDP and GGI has a strong predictive influence on navigation ability, other country-level factors might influence navigation ability. Evidence suggests that driving rather than taking public transport has a positive effect on spatial knowledge [30, 31]. While this might explain why North Americans and Australians are particularly successful as populations compared to equivalent (GDP) European countries that rely more on public transport [32,33], it fails to explain why the Nordic countries perform so well as a group. We speculate that this specificity may be linked to Nordic countries sharing a culture of participating in a sport related to navigation: orienteering. Invented as an official sport in the late 19th century in Sweden, the first orienteering competition open to the public was held in Norway in 1897. Since then, it has been more popular in Nordic countries than anywhere else in the world, and is taught in many schools [34]. We found that ‘orienteering world championship’ country results significantly correlated with countries’ CM (Pearson’s correlation *ρ* = 0.55, *p* = 0.01), even after correcting for GDP per capita (Figure S15). Future targeted research will be required to evaluate the impact of cultural activities on navigation skill.

As mentioned at the beginning of the manuscript, we considered that video games experience might bias performance and therefore we normalized OP with performance on the first two tutorial wayfinding levels, where no navigation skill was required. To further control for gaming skill bias, we re-ran the analyses presented above with a subset of participants with similar performance on the first two tutorial levels and obtained similar results (for more details see Supplementary Discussion and Figure S7b). Another way to control whether Sea Hero Quest captured spatial ability is to quantify how our data fit with results from the spatial cognition literature. For instance, path integration models predict error accumulation over travelled distance and increasing turning angle [35, 36]. We compared this prediction with participants’ performance in the path integration task and showed that in all tested countries, path integration accuracy decreased with complexity (Figure 2F and Figure S10). This shows that real world path integration models do predict performance in Sea Hero Quest path integration task. For more details see the Supplementary Discussion.

## Conclusion

Through the use of an online gaming approach, we have been able to reveal for the first time a global benchmark for spatial navigation. This approach enables predictions to be made about an individual’s spatial navigation performance based on their age, gender and nationality. Specifically, GDP per capita and GGI of countries are predictive of their inhabitants’ average spatial navigation performance. Thus the collected dataset embodies a unique resource, which not only informs our understanding of global cognitive abilities but also provides a stepping stone towards spatial navigation diagnostics and treatment in patient populations with navigation deficits, such as Alzheimer’s pathophysiology [24, 37]. Use of online gaming for the assessment of cognitive abilities has a promising future in particular with an ever-increasing world population taking up gaming as a recreational activity.

### Game Design

To test the global population on their navigation ability we worked with the independent video games design company Glitchers Ltd to produce a video game using Unity 3D (Unity Technologies, Copenhagen Denmark) for smart phones and tablets (apple and android devices). We were supported in this design process by staff at Deutsche Telekom (Germany) and Saatchi and Saatchi London (UK). ‘Sea Hero Quest’ (SHQ) was released on 4 May 2016 on the App Store for iOS and on Google Play for Androids. It is available in 17 languages: English, French, German, Spanish, Macedonian, Greek, Croatian, Dutch, Albanian, Hungarian, Romanian, Slovak, Czech, Polish, Portuguese, Italian, and Serbian, Figure S1a. The game is manipulated through three controls, designed to be intuitive; specifically, these were tap left to turn left, tap right to turn right, swipe up to speed up. Alongside tasks and levels, players were also asked a set of optional questions which included their age, gender and nationality (Figure S4).

### Tasks

The experimental tasks in SHQ were accessed by unlocking levels sequentially (Figure S1c). These levels were comprised of 5 themed areas, each containing 15 levels. Through the game, participants followed a sea captain as he tries to recover his father’s lost memories (Figure S1b). There were three types of task. *Wayfinding levels*: at the beginning of each level, participants were given locations to visit from a map. The map disappeared, and they had to navigate a boat through a virtual environment to find different checkpoints (Figure S2). *Path Integration levels*: participants had to find a flare and shoot it back toward the starting point (Figure S3). *Chase levels*: participants chased a sea creature to take a picture of it. Chase levels were purely for motivational purposes and allow participants the capacity to share their game progress via social media in the form of a ‘photograph’ of the sea creature found. No data was collected from chase levels. Participants are encouraged to collect as many ‘stars’ as possible across the levels: the faster (Wayfinding task) or the more accurate (Path Integration task), the more stars were obtained. These stars unlocked the capacity to modify the boat in the game.

### Participants

Between May 2016 and July 2017, 2,512,123 participants from 255 countries and dependent territories downloaded and completed at least the first level of the game, see Table S1. Amongst them, 1,446,954 (57.6%) entered their age, gender and nationality. Examining the age distribution (Figure S5a), it is evident that age groups 18 and 99 years old contain more participants than would be predicted from the distribution. This is likely due to these numbers being the two extremes of the age range. It is likely that players under the age of 18 may have adopted these age bands. Since it is impossible to separate ‘real’ from ‘fake’ 18 and 99 years old players, for the current analysis we removed these two age groups from the dataset, leaving 926,456 (36.9%) participants. SHQ level progression is linear, i.e. one needs to complete level N in order to unlock level N+1 (Figure S1c). Hence, the number of participants decreases with the progression through the game, as shown in Figure S6. To ensure a good tradeoff between sample size and amount of data per player, we included in the analysis participants who played at least the first 6 wayfinding levels (level numbers 1, 2, 3, 6, 7 and 8) and the first 2 path integration levels (levels 4 and 9). Level 5 is a creature chase level. This represents 625,626 (24.9%) valid participants. To reduce selection bias and ensure stable cross-country comparisons, we only included participants from countries with at least 500 valid participants. As a result of this sampling process, 558,143 (22.2%) participants from 57 countries were included in the analysis. Amongst them, 312,886 males (age: 34.97 ± 14.39 years old) and 245,257 females (age: 35.98 ± 15.50 years old), cf. Table S1 for country by country information.

### Data

Within the opening screen and the Journal menu, participants were made aware of the purpose of the game. They were asked whether they were willing to share their data with us and were guided to where they can opt out. The opt out was always available in the settings. The website for the game (www.seaheroquest.com) was linked to from the About menu and provided full information about the study and what the data was going to be used for. If the participant agreed, their data (boat trajectory flare accuracy and demographics) were anonymously stored in a secure T-Systems server in Germany. The application is managed by T-Systems’ scalable Docker offering called ‘AppAgile’ which is operated out of T-Systems’ datacenter to ensure data integrity and data privacy according to German data security law. The data are owned by Deutsche Telekom and then licensed to University College London for analysis. Each participant was identified by a universally unique identifier (user-uuid), a 128-bit number commonly used to identify information in computer systems. Participant’s sessions were identified by another universally unique identifier (instance-uuid). Only completed levels were stored and analysed. During Wayfinding levels, the coordinates of participants’ trajectories were sampled at Fs = 2 Hz. During Path Integration levels, flare accuracy was quantified in term of stars obtained by the participant. Stars were awarded based on participant’s choice between 3 proposed directions: 3 stars for the correct answer (their starting point), 2 stars for the second closest direction, and 1 star for the third closest direction.

### Metrics

To quantify spatial abilities, three measures were computed for each level.

- Trajectory length in pixels, defined as the Euclidean distance 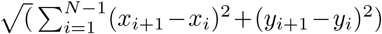 with (*x*_*i*_, *y*_*i*_)_*i*∈[1‥*N*]_ a N-points trajectory.
- Duration to complete the game in seconds. Since *F*_*s*_ = 2 Hz, Duration = *N*/2).
- Flare accuracy in number of stars: 1, 2 or 3.

To correct for video gaming skill, we normalized durations and trajectory lengths by dividing them by the sum of their values at the first two levels, where no sense of direction is needed (Supplementary Discussion).

We found all three measures correlated: Pearson’s correlation between trajectory length and duration *r* = 0.75, *p* < 0.001; trajectory length and path integration accuracy *r* = −0.21, *p* < 0.001; duration and path integration accuracy *r* = −0.20, *p* < 0.001. While path integration relied on the integration of perceived ego motion information over time, wayfinding required the planning of multi-stop routes, memory of survey/map, transformation of survey representation into egocentric reference frame. It is therefore not surprising that trajectory lengths and duration - both wayfinding-related metrics - were more correlated with each other than with flare accuracy.

We defined an overall performance metric (OP) summarizing normalized durations, trajectory lengths and flare accuracies. OP was the 1st component of a Principal Component Analysis across the normalized durations and trajectory lengths of levels 6, 7, 8 and the flare accuracies of levels 4 and 9 (66.9% of the variance explained). OP captured different aspects of navigation abilities, providing a more general performance measure, which reflects overall navigation abilities covering a range of tasks.

### Data Availability

Given the magnitude of the dataset, the authors made it available from a dedicated server. Access will be granted upon reasonable request.

### Code Availability

Analyses and figures were made using Python, JavaScript, R, and Matlab. Scripts are available from the corresponding author upon request.

### Ethics

This study has been approved by UCL Ethics Research Committee. The ethics project ID number is CPB/2013/015

## Acknowledgments

The authors wish to thank Deutsche Telekom for supporting and funding this research, Alzheimer Research UK for funding the analysis, the Glitchers Limited for the game production, Saatchi and Saatchi London for project management and creative input, Dara Mohammadi and Rogier Kievit for their advice.

## Author Contributions

HS and MH supervised the project, HS, MH, SS, JW, and CH designed research; AC, RS, WdC, and EM analyzed data; AC, MH and HS wrote the paper.

## References

1. Dehaene, S., Izard, V., Spelke, E. & Pica, P. Log or Linear? Distinct Intuitions of the Number Scale in Western and Amazonian Indigene Cultures. Science 320, 1217–1220 (2008).

2. Henrich, J., Heine, S. J. & Norenzayan, A. The weirdest people in the world? Behavioral and Brain Sciences 33, 61–135 (2010).

3. Chiu, M. M. & Klassen, R. M. Relations of mathematics self-concept and its calibration with mathematics achievement: Cultural differences among fifteen-year-olds in 34 countries. Learning and Instruction 20, 2–17 (2010).

4. Hanushek, E. A. & Woessmann, L. Do better schools lead to more growth? cognitive skills, economic outcomes, and causation. Journal of Economic Growth 17, 267–321 (2012).

5. Haun, D. B. M., Rapold, C. J., Janzen, G. & Levinson, S. C. Plasticity of human spatial cognition: Spatial language and cognition covary across cultures. Cognition 119, 70–80 (2011).

6. Lovett, A. & Forbus, K. Cultural commonalities and differences in spatial problem-solving: A computational analysis. Cognition 121, 281–287 (2011).

7. Goeke, C. et al. Cultural background shapes spatial reference frame proclivity. Scientific Reports 5, 1–13 (2015).

8. Darwin, C. R. Origin of Certain Instincts. Nature 7, 417–418 (1873).

9. Wiener, J. M., Büchner, S. J. & Hölscher, C. Taxonomy of human wayfinding tasks: A knowledge-based approach. Spatial Cognition & Computation 9, 152–165 (2009).

10. Mittelstaedt, M.-L. & Mittelstaedt, H. Homing by path integration in a mammal. Naturwissenschaften 67, 566–567 (1980).

11. Etienne, A. S. & Jeffery, K. J. Path integration in mammals. Hippocampus 14, 180–192 (2004).

12. Baker, D. P. & Jones, D. P. Creating gender equality: Cross-national gender stratification and mathematical performance. Sociology of Education 66, 91–103 (1993).

13. Else-Quest, N. M., Hyde, J. S. & Linn, M. C. Cross-national patterns of gender differences in mathematics: A meta-analysis. Psychological Bulletin 136, 103–127 (2010).

14. Guiso, L., Monte, F., Sapienza, P. & Zingales, L. Culture, Gender, and Math. Science 320, 1164–1165 (2008).

15. Hyde, J. S. & Mertz, J. E. Gender, culture, and mathematics performance. Proceedings of the National Academy of Sciences 106, 8801–8807 (2009).

16. Reilly, D. Gender, Culture, and Sex-Typed Cognitive Abilities. PLoS ONE 7, e39904–16 (2012).

17. Lippa, R. A., Collaer, M. L. & Peters, M. Sex Differences in Mental Rotation and Line Angle Judgments Are Positively Associated with Gender Equality and Economic Development Across 53 Nations. Archives of Sexual Behavior 39, 990–997 (2010).

18. Ghisletta, P., Rabbitt, P., Lunn, M. & Lindenberger, U. Two thirds of the age-based changes in fluid and crystallized intelligence, perceptual speed, and memory in adulthood are shared. Intelligence 40, 260–268 (2012).

19. Anguera, J. A. et al. Video game training enhances cognitive control in older adults. Nature 501, 97–101 (2013).

20. Lindenberger, U. Human cognitive aging: Corriger la fortune? Science 346, 572–578 (2014).

21. Lindenberger, U., Singer, T. & Baltes, P. B. Longitudinal Selectivity in Aging Populations: Separating Mortality-Associated Versus Experimental Components in the Berlin Aging Study (BASE). Journal of Gerontology: Psychological Sciences 57B, 474–482 (2002).

22. Moffat, S. D. Aging and Spatial Navigation: What Do We Know and Where Do We Go? Neuropsychology Review 19, 478–489 (2009).

23. Klencklen, G., Després, O. & Dufour, A. What do we know about aging and spatial cognition? Reviews and perspectives. Ageing Research Reviews 11, 123–135 (2012).

24. Lester, A. W., Moffat, S. D., Wiener, J. M., Barnes, C. A. & Wolbers, T. The aging navigational system. Neuron 95, 1019–1035 (2017).

25. Wang, H. & Song, M. Ckmeans. 1d. dp: optimal k-means clustering in one dimension by dynamic programming. The R Journal 3, 29 (2011).

26. Skirbekk, V. & Loichinger, E. Variation in cognitive functioning as a refined approach to comparing aging across countries. Proceedings of the National Academy of Sciences 109 (2012).

27. Gunderson, E. A., Ramirez, G., Beilock, S. L. & Levine, S. C. The relation between spatial skill and early number knowledge: the role of the linear number line. Developmental Psychology 48, 1229 (2012).

28. Uttal, D. H., Meadow, N. G., Hand, E. T. L. L., Warren, C. & Newcombe, N. S. The malleability of spatial skills: A meta-analysis of training studies. Psychological Bulletin 139, 352–402 (2013).

29. Poumanyvong, P., Kaneko, S. & Dhakal, S. Impacts of urbanization on national transport and road energy use: Evidence from low, middle and high income countries. Energy Policy 46, 268–277 (2012).

30. Maguire, E. A., Woollett, K. & Spiers, H. J. London taxi drivers and bus drivers: a structural mri and neuropsychological analysis. Hippocampus 16, 1091–1101 (2006).

31. Sandamas, G. & Foreman, N. Active Versus Passive Acquisition of Spatial Knowledge While Controlling a Vehicle in a Virtual Urban Space in Drivers and Non-Drivers. SAGE Open 5, 1–9 (2015).

32. Pucher, J. Urban travel behavior as the outcome of public policy: the example of modal-split in western europe and north america. Journal of the American Planning Association 54, 509–520 (1988).

33. Bassett, D. R., Pucher Jr, J., Buehler, R., Thompson, D. L. & Crouter, S. E. Walking, cycling, and obesity rates in europe, north america, and australia. Journal of Physical Activity and Health 5, 795–814 (2008).

34. Annerstedt, C. Physical education in scandinavia with a focus on sweden: a comparative perspective. Physical Education and Sport Pedagogy 13, 303–318 (2008).

35. Benhamou, S. & Séguinot, V. How to find one’s way in the labyrinth of path integration models. Journal of Theoretical Biology 174, 463–466 (1995).

36. Wiener, J. M., Berthoz, A. & Wolbers, T. Dissociable cognitive mechanisms underlying human path integration. Experimental Brain Research 208, 61–71 (2011).

37. Kunz, L. et al. Reduced grid-cell–like representations in adults at genetic risk for alzheimer’s disease. Science 350, 430–433 (2015).

